# Data mining patented antibody sequences

**DOI:** 10.1101/2020.11.26.389866

**Authors:** Konrad Krawczyk, Andrew Buchanan, Paolo Marcatili

## Abstract

Patent literature should be a reflection of thirty years of engineering efforts in developing monoclonal antibody therapeutics. Such information is potentially valuable for rational antibody design. Patents however are not designed to convey scientific knowledge, but rather legal protection. It is unclear whether antibody information from patent documents, such as antibody sequences could be useful for the therapeutic antibody sphere in conveying engineering know-how rather than act as legal reference only. To assess the utility of patent data for therapeutic antibody engineering, we quantified the amount of antibody sequences in patents destined for medicinal purposes and how well they reflect the primary sequences of therapeutic antibodies in clinical use. We identified 16,526 patent families from major jurisdictions (e.g. USPTO and WIPO) that contained antibody sequences. These families held 245,109 unique antibody chains (135,397 heavy chains and 109,712 light chains) that we compiled in our Patented Antibody Database (PAD, http://naturalantibody.com/pad). We find that antibodies make up a non-trivial proportion of all patent amino acid sequence depositions (e.g. 10.95% of USPTO Full Text database). Our analysis of the 16,526 families demonstrates that the volume of patent documents with antibody sequences is growing with the majority of documents classified as containing antibodies for medicinal purposes. We further studied the 245,109 antibody chains from patent literature to reveal that they very well reflect the primary sequences of antibody therapeutics in clinical use. This suggests that patent literature could serve as a reference of previous engineering efforts to improve rational antibody design.

## Introduction

The binding versatility of antibodies has been used for medicinal purposes making them the most successful group of biotherapeutics^1^. Typical timelines involved in bringing these molecules to the market are slow, however more and more molecules are approved in the US and EU each year^1^. Successful exploitation of antibodies by either experimental^2,3^ or computational techniques^4^ relies on our ability to understand what makes a successful antibody-based therapeutic^5,6^.

Therapeutic antibodies on the market and in late stage clinical trials have been previously studied by experimental^2,7^ and computational^6^ approaches to identify properties that make a successful biotherapeutic. Such studies^2^ however only focused on 137 approved or post-phase-I antibodies (Clinical Stage Therapeutics, or CSTs), which is a small dataset in the light of the mutational space available to antibodies^8^. CSTs, are high-quality data-points that are end results of a long engineering process of selecting a molecule from a number of viable candidates. The single successful therapeutic molecule is therefore only partially representative of the engineering process. Full public disclosure of the efforts involved in developing a therapeutic antibody constituting intermediate sequences and selection decisions is not desirable because of the commercial value of such know-how, which needs to be legally protected.

Because of the need to protect the know-how involved in engineering therapeutic antibodies, relevant information needs to be disclosed in patent documents. Previous approaches to extract information on patent antibody landscape^9^ or specific antibody formats^10^ focused on keyword and patent classification searches. One can broadly discern between patents on antibody techniques (e.g. phage display, humanization) and novel antibody molecules. It is the patents on novel molecules that could be of particular engineering interest as these reflect the constructs that might find their way into the clinic. The disclosure of antibody sequence and target information^11^ in such patents reveals to a certain extent the engineering choices as such molecules have been subjected to myriad prior tests to be suitable candidates for expensive legal protection and further clinical trials.

The purpose of patent literature is not conveying scientific knowledge, but legal protection. In this work we assessed the utility of patent data for therapeutic antibody engineering efforts by establishing the extent to which antibodies from patents reflect therapeutics in clinical use. For this purpose, we identified patent documents that contained antibody sequences, to quantify how many of these were destined for medicinal purposes and how well they reflect advanced stage therapeutics.

## Results

### Antibodies account for a non-trivial proportion of sequences deposited in patent documents

We identified documents with antibody sequences by downloading data from four data sources: USPTO (http://uspto.gov), WIPO (http://wipo.int), DDBJ^12^, and EBI^13^. Choice of the data sources was motivated by the availability of biological sequences and coverage of patent documents worldwide. Biological sequence information is not universally available in patent documents in all jurisdictions^14^. In certain cases, the data is not freely available, but rather accessible for a fee (e.g. European Patent Office). Primary access to biological sequences in machine-readable format is freely available from the USPTO and WIPO. USA is the largest pharmaceutical market^15^, compelling pharmaceutical companies developing a novel antibody therapeutic to seek patent protection within the jurisdiction of USPTO. Similarly, it is common to seek protection under the auspices of WIPO PCT system in order to spread the coverage of the patent documents across many jurisdictions worldwide. Furthermore, data from certain major jurisdictions, such as EPO, JPO and KPO are available via third parties such as DDBJ^12^ and EBI^13^. Therefore, we argue that datasets made available via USPTO, DDBJ, WIPO and EBI provide a reasonable coverage of the worldwide antibody sequence patents.

We extracted raw sequence data from USPTO, WIPO, DDBJ and EBI on Jan 30^th^ 2020, with the particulars of parsing the heterogenous sources described in Methods. From each dataset we extracted raw, redundant amino acid and nucleic acids sequences. Sequences containing exclusively nucleotides were translated to amino acids using IgBlast^16^ as described previously^17^. Raw amino acid sequences were analyzed using ANARCI^18^ to identify antibody variable region chains (VH, VL, including scFvs). We report the number of raw sequences analyzed and the resulting identified antibodies in Table 1.

**Table 1.** Published biological sequences and proportion thereof identified as antibody chains. We extracted raw sequences from USPTO (divided between the full text, FT, and long listing repository PSIPS), DDBJ, WIPO and EBI. The total number of raw sequences is given in column Total Raw. Of these we show how many were identified by ANARCI as containing an antibody chain (column Ab-identified). In the column “% Total” we report the proportion of identified antibody sequences out of the total of raw sequences. Both Total Raw and Ab-identified columns report the redundant number of sequences so as to exemplify the volume of antibody depositions in patent sequences – we report the number of unique heavy (H) and light (L) chains in the parentheses in column “Ab-identified”.

| Source | Sequence Type | Total Raw | Ab-identified<br>(unique Heavy (H), Light (L)) | %Total |
| --- | --- | --- | --- | --- |
| USPTO FT | Amino Acid | 5,534,127 | 606,036<br>(H=52,388,L=38,922) | 10.95 |
|  | Nucleotide | 7,068,248 | 229,547<br>(H=21,169,L=17,009) | 3.24 |
| USPTO PSIPS | Amino Acid | 25,527,942 | 470,317<br>(H=33,806,L=24,086) | 1.84 |
|  | Nucleotide | 176,840,912 | 376,567<br>(H=35,802,L=46,374) | 0.21 |
| DDBJ | Amino Acid | 4,412,209 | 533,762<br>(H=61,999,L=46,015) | 12.09 |
|  | Nucleotide | 44,968,142 | 413,485<br>(H=35,290,L=28,502) | 0.91 |
| WIPO | Amino Acid | 10,275,174 | 435,218<br>(H=67,533,L=49,699) | 4.23 |
|  | Nucleotide | 13,490,560 | 160,542<br>(H=35,747,L=27,275) | 1.19 |
| EBI | Amino Acid | 10,368,431 | 713,620<br>(H=73,450,L=50,326) | 6.88 |
|  | Nucleotide | 12,349,772 | 38,366<br>(H=15,792,L=13,339) | 0.31 |

We find a higher proportion of sequences identified as antibodies in amino acid depositions which account for as many as 10.95% and 12.09% of USPTO-FT and DDBJ datasets respectively. In fact, large portion of sequences deposited in patents are very short; for instance in USPTO-FT only 1,811,694 (32.73%) amino acid sequences are longer than 50 amino acids, and antibodies make up 30.50% of these. This stands to show that antibodies make up a non-trivial volume of all the sequences deposited in patent documents.

Antibody sequence data in patents is however redundant to a large extent when one considers a unique sequence to be defined by its variable region. Combining all the non-redundant VH and VL sequences from our datasets we count 245,109 unique antibody domains (135,397 heavy chains and 109,712 light chains). This suggests that many antibody variable region sequences are listed as part of multiple patent documents. Not all of these sequences however are guaranteed to have been developed for medical applications, which can be determined by analyzing the text content of patent documents.

### Patent landscape of documents containing antibody sequences

We analyzed the text content of patents containing antibody sequences so as to establish what proportion of these list molecules for medicinal purposes. We connected all the redundant antibody sequences to their patent documents and identified a patent family for each. A patent family can be regarded as identifying documents with the same subject matter across several jurisdictions. Altogether our 245,109 sequences are distributed among 16,526 patent families. We extracted the metadata from the patent documents, such as titles, abstracts, inventors and classifications. We used this information to determine the proportion of patents destined for medicinal applications by analyzing their classifications, and whether the inventors and listed targets resemble entities and molecules associated with development of monoclonal antibody therapies.

### Most patent documents citing antibody sequences are destined for medicinal applications

We analyzed the patent classifications of the 16,526 patent families that indicate the purpose of the invention described in each document. We extracted the Cooperative Patent Classification (CPC, developed by USPTO and EPO, https://www.cooperativepatentclassification.org/) designations from the documents as this was the most common listed scheme, covering 15,951 (96.52%) out of 16,526 families. Patent classifications according to CPC have a section, class, subgroup, main group and a subgroup (e.g. classification C07K16/2866 has section C, class 07, subclass K, main group 16 and subgroup 2866). We divided the 15,951 families according to their CPC classifications excluding the subgroup (e.g. C07K16/2866 becomes C07K16) to reveal the general categories the documents fall into and present results in Table 2.

**Table 2.** Subclasses of the patent classifications. Most common subclasses associated with patents including antibody sequences according to the Cooperative Patent Classification (CPC, https://www.cooperativepatentclassification.org/). There were 15,951 patents containing antibodies with CPC classification and the percentage of families in each class is expressed as a proportion of this number.

| CLASS | TOTAL<br>FAMILIES (%) | DESCRIPTION |
| --- | --- | --- |
| **C07K16** | 13,790 (86.4) | Immunoglobulins [IGs], e.g. monoclonal or polyclonal antibodies (antibodies with enzymatic activity, e.g. abzymes) |
| **C07K2317** | 12,001 (75.2) | Immunoglobulins specific features |
| **A61K39** | 9,459 (59.3) | Medicinal preparations containing antigens or antibodies |
| **C07K2319** | 3,451 (21.6) | Fusion polypeptide |
| **G01N33** | 3,105 (19.4) | Investigating or analysing materials by specific methods not covered by groups G01N1/00 - G01N31/00 |
| **C07K14** | 3,037 (19.0) | Peptides having more than 20 amino acids; Gastrins; Somatostatins; Melanotropins; Derivatives thereof |
| **A61K47** | 2,392 (14.9) | Medicinal preparations characterised by the non-active ingredients used, e.g. carriers or inert additives; Targeting or modifying agents chemically bound to the active ingredient |
| **A61K38** | 2,058 (12.9) | Medicinal preparations containing peptides |
| **C12N15** | 1,972 (12.3) | Mutation or genetic engineering; DNA or RNA concerning genetic engineering, vectors, e.g. plasmids, or their isolation, preparation or purification; Use of hosts therefor |
| **A61P35** | 1,900 (11.9) | Specific therapeutic activity of chemical compounds or medicinal preparations |
| **A61K45** | 1,671 (10.4) | Medicinal preparations containing active ingredients not provided for in groups |
| **A61K31** | 1,415 (8.8) | Medicinal preparations containing organic active ingredients |
| **G01N2333** | 1,329 (8.3) | Assays involving biological materials from specific organisms or of a specific nature |
| **C12N5** | 816 (5.1) | Undifferentiated human, animal or plant cells, e.g. cell lines; Tissues; Cultivation or maintenance thereof; Culture media therefor |

Subgroup C07K16 that indicates immunoglobulins, is the most common classification, present in 13,790 (86.45%) of the 15,951 patent families. Families listing antibodies for medicinal purposes (A61K39) account for 9,459 (59.30%) of the 15,951 families. Furthermore the more general medicinal categorization A61K (preparations for medical, dental or toilet purposes) accounts for 11,398 (71.45%) of the 15,951 patent families. This indicates that the majority of documents citing antibody sequences are developed for medicinal purposes, such as novel treatments or diagnostics. This is well reflected by the organizations that submit such patent applications, where 9 out of top 10 and 69 out of top 100 are pharmaceutical companies associated with development of monoclonal antibody therapies for a range of targets and disease indications (see Supplementary Section 1).

### Targets of antibodies in patent documents correspond to known therapeutic targets

We checked to what extent antibody targets reported in patent literature reflect those of known therapeutic antibodies. Each patent family in PAD was scanned for antibody target (see Methods). Therapeutic antibodies in clinical use together with their associated targets were compiled from the WHO lists of International Nonproprietary Names^19^ (INNs, e.g. list 122^20^) IMGT ^21^, Antibody Society (http://www.antibodysociety.org) and Thera-SAbDab^22,23^, resulting in 563 unique INNs. We grouped the targets by number of patent families and therapeutic antibodies they were associated with. We present the results for top 30 targets sorted by the highest number of patent families in Table 3.

**Table 3.** Top 30 targets in patent documents. We extracted the targets of the antibodies in patent documents and present top 30 ranked by the number of families where they were mentioned. For each target, we show the number of patent families mentioning the target (#Families), the number of therapeutics on the market/in the clinic against it (#Therapeutics) and the cumulative number of therapeutics covered by the top targets (#Therapeutics cumulative).

| RANK | TARGET | #FAMILIES | #THERAPEUTICS | #THERAPEUTICS<br>(CUMULATIVE) |
| --- | --- | --- | --- | --- |
| 1 | pd1 | 284 | 20 | 20 |
| 2 | cd3 | 221 | 20 | 40 |
| 3 | her1 | 190 | 17 | 57 |
| 4 | pdl1 | 189 | 12 | 69 |
| 5 | tnfa | 185 | 6 | 75 |
| 6 | her2 | 175 | 9 | 84 |
| 7 | cd20 | 169 | 14 | 98 |
| 8 | influenza | 151 | 5 | 103 |
| 9 | cmet | 136 | 4 | 107 |
| 10 | vegfa | 135 | 7 | 114 |
| 11 | amyloid beta | 129 | 8 | 122 |
| 12 | hiv | 123 | 2 | 124 |
| =13 | il6 | 102 | 7 | 131 |
| =13 | cd40 | 102 | 9 | 140 |
| =13 | cd19 | 102 | 6 | 146 |
| 14 | ctla4 | 97 | 6 | 152 |
| =15 | il17 | 92 | 8 | 160 |
| =15 | igf1r | 92 | 6 | 166 |
| 16 | pcsk9 | 89 | 8 | 174 |
| 17 | her3 | 87 | 6 | 180 |
| =18 | rsv | 73 | 5 | 185 |
| =18 | cd38 | 73 | 5 | 190 |
| =19 | tau | 68 | 5 | 195 |
| =19 | lag3 | 68 | 7 | 202 |
| 20 | ox40 | 65 | 6 | 208 |
| 21 | bcma | 64 | 1 | 209 |
| =22 | il23 | 63 | 5 | 214 |
| =22 | cd47 | 63 | 2 | 216 |
| 23 | ang2 | 62 | 2 | 218 |
| 24 | vegfr2 | 56 | 5 | 223 |

The number of patent families associated with a target appears to correspond to a larger number of therapeutic antibodies against the same target. Top 10 targets sorted by number of their patent families account for 114 (20.24%), top 30 account for 223 (39.60%) and top 100 account for 369 (65.54%) out of 563 therapeutics. Therefore, targets from patents listing antibody sequences provide a reasonable reflection of the targets of currently available therapeutic antibodies. In fact the greater number of patent families per target can be associated with an earlier date of the said target being mentioned in a patent document (Figure 1A). It does not mean however that the patent space for monoclonal antibodies is saturated as the number of new targets mentioned is increasing (Figure 1B). This suggests that studying patent documents including antibody sequences could provide an early indication of their targets and thus activity in the field of therapeutic antibodies.

**Figure 1.**
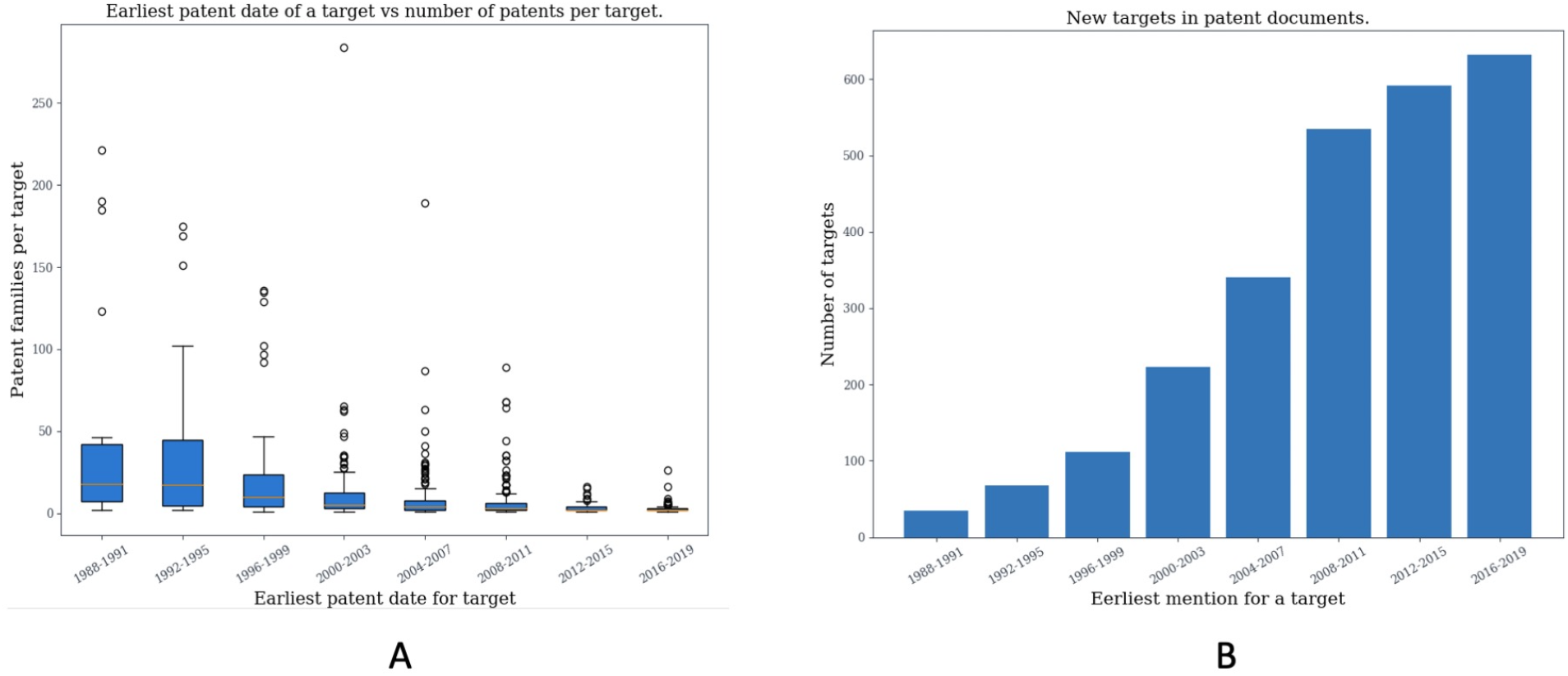
Target usage in patent documents reporting antibody sequences. A) Relationship between number of patent families per target and the earliest mention of the target in patent documents containing antibodies. For each target, we noted the earliest date among patent documents citing it and grouped these into 4-year intervals. Within each interval we noted the total number of patent families for a given target and plotted the aggregate for each time interval. B) For each 4-year interval, we plot the number of new target names that were first introduced in a patent document at that time.

### There is a growing number of patent documents associated with antibody sequences

We analyzed the timestamps associated with patents in order to check whether there is a growing trend in releasing documents with antibody sequences and what proportion thereof is made up of molecules for medicinal indications. Each patent family lists several dates corresponding to the activity associated with the patent. We noted the earliest and most recent dates for each patent family to reflect the original submission dates and the most up-to-date activity respectively.

We plotted the earliest dates for each patent family in our dataset which indicates that the number of patent documents containing antibody sequences is steadily rising (Figure 2). The most recent dates associated with the same patent documents (Figure 2) shows a more acute rise since 2016 which indicates strong activity within the earlier submitted patents. Since not all patent families are explicitly destined for medicinal applications, we have plotted the corresponding earliest and most recent dates for the 9,459 documents classified as medicinal preparations containing antibodies (Supplementary Figure 1) which recapitulates the increasing number of patent documents being released.

**Figure 2.**
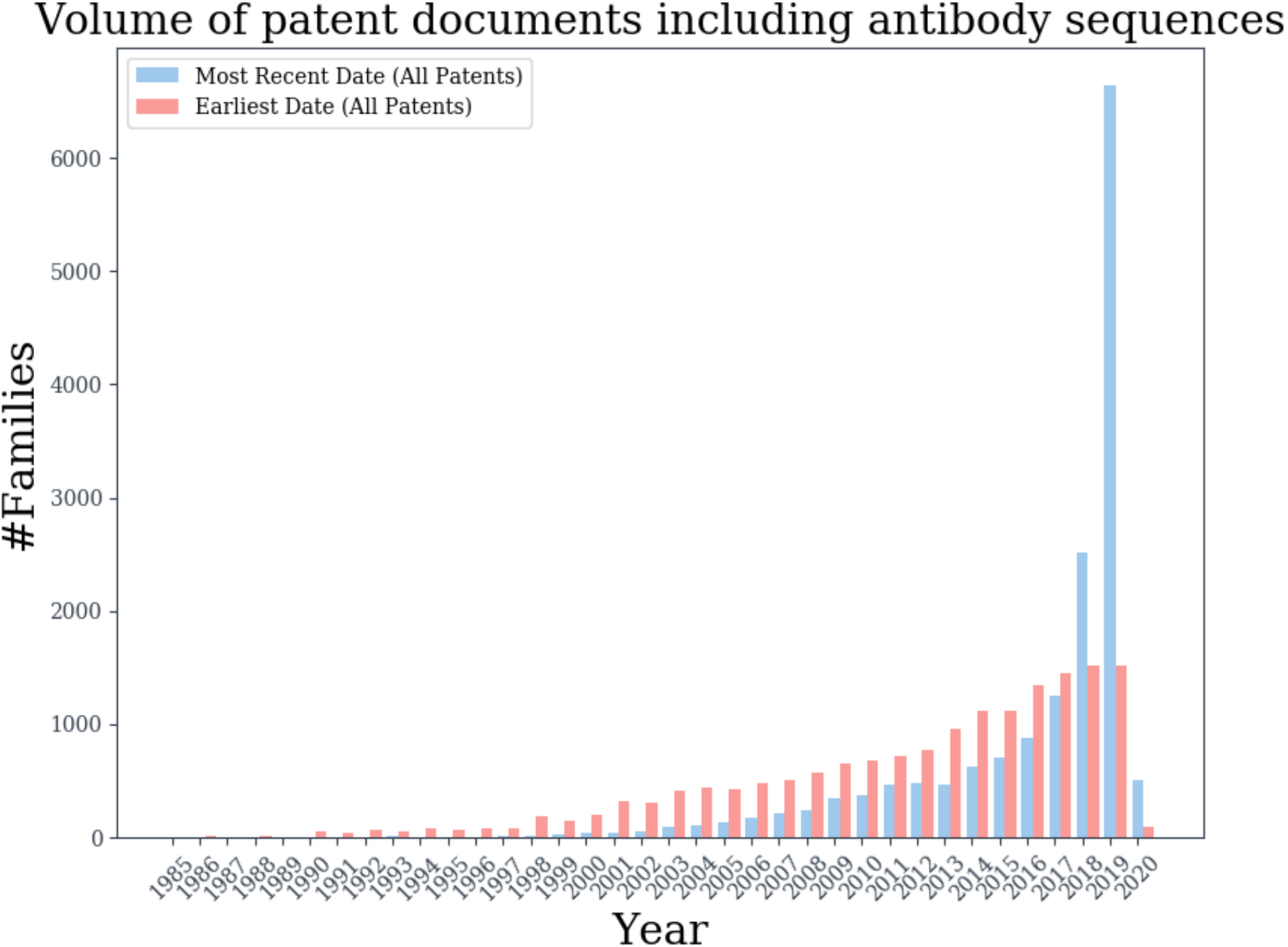
The volume of patent family documents listing antibody sequences per year. For each patent family we noted the earliest and most recent dates of any documents associated with it and the aggregate numbers of these are given by red and blue bars respectively. The apparent low activity in 2020 can be attributed to the fact that data contributed in 2020 only account for January that year.

Increasing patent activity in documents listing antibody sequences for medicinal indications is in line with the rising approval rates for antibody-based biologics^1,24^. Given that the patents are an early sign of approvals to come, it suggests that we can expect more biologics in the clinics in the foreseeable future. Since majority of such patent documents are indeed listing antibodies for medicinal purposes, the broad characteristics of the molecules listed in patent documents could provide an indication of the engineering choices in their design.

### The sequence landscape of patented antibodies

Antibody sequences found in patent documents could reflect the broad decisions taken by engineers shaping these molecules before they arrive in the clinic. However, not all antibody sequences found in patent documents are destined for medicinal applications. For this reason we analyzed the broad sequence characteristics of antibodies from patent documents to establish to what extent they are a reflection of therapeutic antibodies in clinical use and vice-versa. We performed this analysis by looking at all of our antibodies from all patents (AllPatAb) and just the subset associated with documents classified as containing antibodies for medicinal applications (MedPatAb). Altogether AllPatAb consisted of 135,397 heavy chains and 109,712 light chains whereas MedPatAb consisted of 93,067 heavy chains (68.73% of all heavy chains) and 67,667 light chains (67.67% of all light chains).

### Most antibody sequences from patents align to human and mouse germline V region genes

We checked the patterns of organism-specific germline gene usage in antibody sequences originating from patent documents. Since organism reporting is not consistent in patent documents, we aligned the sequences in PAD to HMMs created from IMGT germline sequences for fifteen organisms: human, mouse, alpaca, rhesus, rabbit, rat, pig, cow, macaque, zebrafish, trout, salmon, dog, horse and chicken. For each organism and germline, we noted the total number of patent antibody sequences aligning to a given germline as well as the number of families they originated from.

We show the number of MedPatAb sequences that aligned to one of our fifteen organisms in Table 4 with the corresponding distribution for AllPatAb sequences in supplementary Table 2. Majority of the unique heavy sequences from patents for medicinal indications align to human germlines (72.80% of unique sequences), followed by mouse (15.39% of unique sequences). The same holds true for light chains with 67.72% of MedPatAb sequences aligning to human and 19.68% to mouse germlines. Antibodies aligning to either mouse or human germlines are most frequently found within protein families. Human-aligned heavy and light chains can be identified in 75.69% and 69.76% patent families respectively. Mouse-aligned heavy and light chains can be found in 52.18% and 53.93% patent families respectively. This broad proportion is also reflected in all the antibody sequences from patents (AllPatAb), indicating that the medicinal patent classification does not skew the broad trend of majority of patented sequences aligning to human and mouse germlines. The alignment to those two organisms reasonably reflects the human focus of antibody development and the rodent antibodies that often server as a basis for humanized therapeutics^25^.

**Table 4.** Most common V-region gene species antibodies from patents aligned to. Antibodies from patent documents destined for medicinal indications (MedPatAb) were aligned to fifteen IMGT-derived^28^ V region germlines from human, mouse, alpaca, rhesus, rabbit, rat, pig, cow, macaque, zebrafish, trout, salmon, dog, horse and chicken. We noted the number of patent sequences that aligned to the given species germline (#Unique Sequences) and the number of patent families (#Patent Families) these originated from.

| HEAVY CHAIN | PER SEQUENCE |  |  | PER FAMILY |  |  |
| --- | --- | --- | --- | --- | --- | --- |
|  | Organism | #Unique Sequences | Percentage | Organism | #Patent Families | Percentage |
|  | human | 67754 | 72.80 | human | 7070 | 75.69 |
|  | mouse | 14326 | 15.39 | mouse | 4874 | 52.18 |
|  | alpaca | 7047 | 7.57 | macaque | 485 | 5.19 |
|  | rabbit | 1313 | 1.41 | horse | 473 | 5.06 |
|  | macaque | 1035 | 1.11 | alpaca | 403 | 4.31 |
|  | horse | 799 | 0.85 | rabbit | 256 | 2.74 |
|  | chicken | 417 | 0.44 | chicken | 46 | 0.49 |
|  | dog | 291 | 0.31 | dog | 28 | 0.29 |
|  | rhesus | 43 | 0.04 | rhesus | 23 | 0.24 |
|  | cow | 30 | 0.03 | cow | 11 | 0.11 |
|  | pig | 9 | ~0 | pig | 9 | 0.09 |
|  | rat | 2 | ~0 | rat | 3 | 0.03 |
|  | salmon | 1 | ~0 | salmon | 2 | 0.02 |
| LIGHT CHAIN | Organism | #Unique Sequences | Percentage | Organism | #Families | Percentage |
|  | human | 45828 | 67.72 | human | 6312 | 69.76 |
|  | mouse | 13320 | 19.68 | mouse | 4880 | 53.93 |
|  | rhesus | 5333 | 7.88 | rhesus | 2238 | 24.73 |
|  | rabbit | 1438 | 2.12 | rat | 361 | 3.98 |
|  | rat | 778 | 1.14 | rabbit | 240 | 2.65 |
|  | chicken | 505 | 0.74 | chicken | 46 | 0.5 |
|  | dog | 220 | 0.32 | cow | 31 | 0.34 |
|  | cow | 213 | 0.31 | dog | 21 | 0.23 |
|  | pig | 17 | 0.02 | pig | 7 | 0.07 |
|  | horse | 15 | 0.02 | horse | 7 | 0.07 |

### Germline V gene usage of antibodies from patent documents corresponds to a large extent with germline V gene usage of therapeutic antibodies

Given that majority of antibodies from patents align to human germlines, we stratified these by the particular human V-region genes. In Table 4 and 5 we report the most common V-region genes medicinal patent sequences align to (corresponding numbers for all patents can be found in Supplementary Table 3). We compare the distribution of germline genes in patents to the germline usage in therapeutic antibodies to show to what extent patent submissions reflect current therapeutics.

**Table 5.**
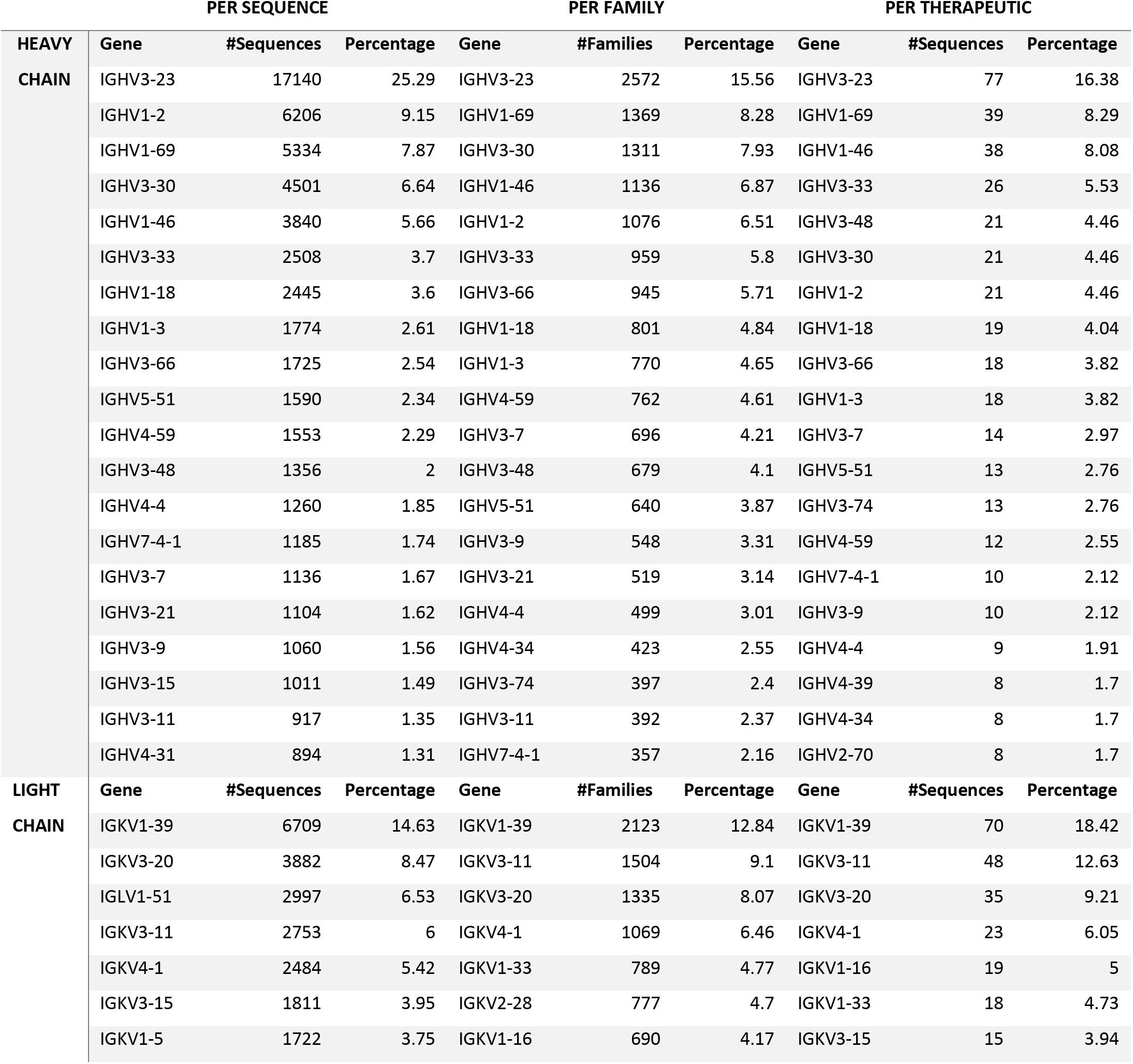
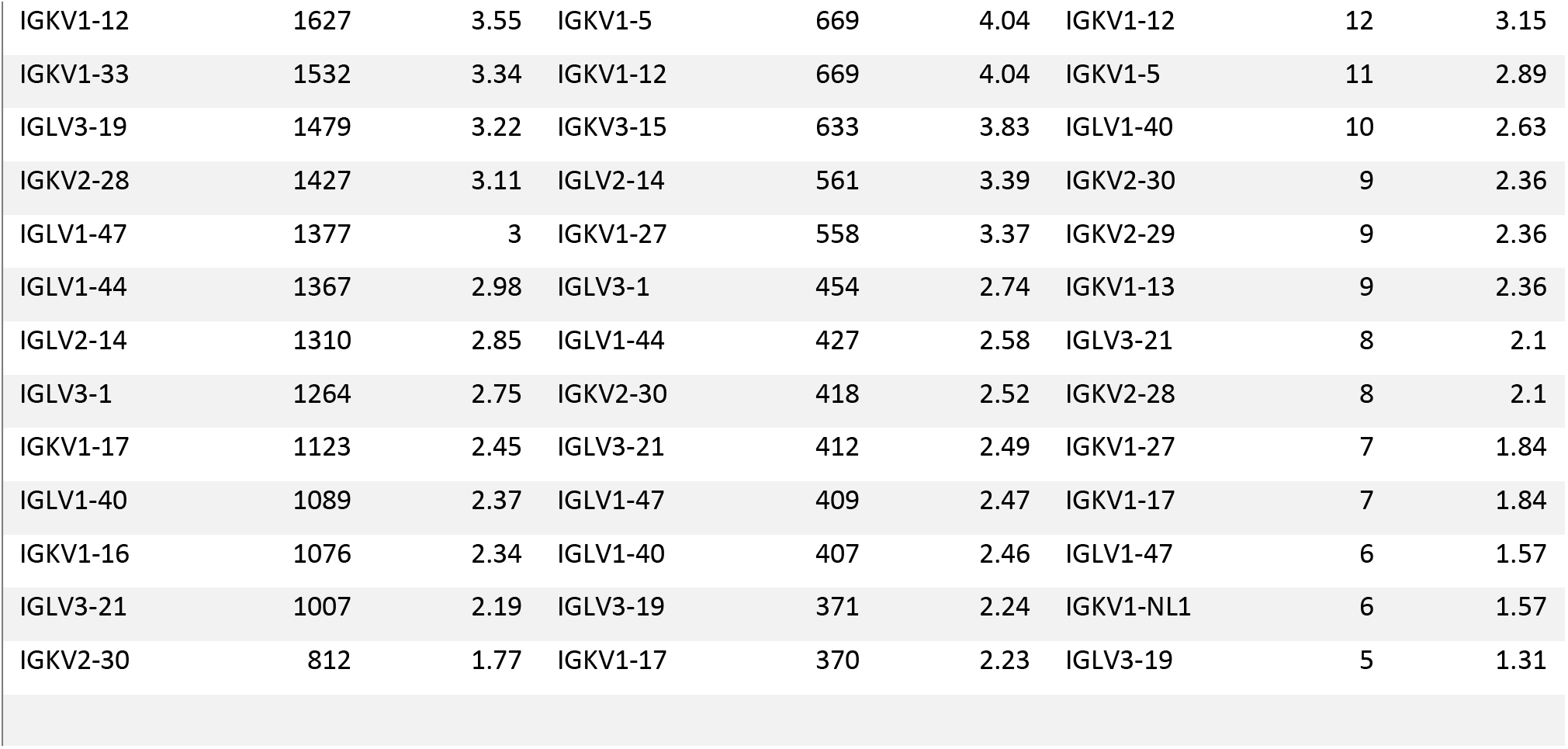
Top-20 most common human V-region genes antibodies from patents aligned to. For each patent antibody sequence for medicinal applications (MedPatAb) that aligned to human germline V-regions, we noted the IMGT V-region gene. We show the number of unique sequences that aligned to a given human V-region gene (Per Sequence) and number of patent families these originated from (Per Family). We also show the number of therapeutic antibody sequences in clinical use that align to the given V-region gene (Per Therapeutic).

| HEAVY CHAIN | PER SEQUENCE |  |  | PER FAMILY |  |  | PER THERAPEUTIC |  |  |
| --- | --- | --- | --- | --- | --- | --- | --- | --- | --- |
|  | Gene | #Sequences | Percentage | Gene | #Families | Percentage | Gene | #Sequences | Percentage |
| HEAVY CHAIN | IGHV3-23 | 17140 | 25.29 | IGHV3-23 | 2572 | 15.56 | IGHV3-23 | 77 | 16.38 |
|  | IGHV1-2 | 6206 | 9.15 | IGHV1-69 | 1369 | 8.28 | IGHV1-69 | 39 | 8.29 |
|  | IGHV1-69 | 5334 | 7.87 | IGHV3-30 | 1311 | 7.93 | IGHV1-46 | 38 | 8.08 |
|  | IGHV3-30 | 4501 | 6.64 | IGHV1-46 | 1136 | 6.87 | IGHV3-33 | 26 | 5.53 |
|  | IGHV1-46 | 3840 | 5.66 | IGHV1-2 | 1076 | 6.51 | IGHV3-48 | 21 | 4.46 |
|  | IGHV3-33 | 2508 | 3.7 | IGHV3-33 | 959 | 5.8 | IGHV3-30 | 21 | 4.46 |
|  | IGHV1-18 | 2445 | 3.6 | IGHV3-66 | 945 | 5.71 | IGHV1-2 | 21 | 4.46 |
|  | IGHV1-3 | 1774 | 2.61 | IGHV1-18 | 801 | 4.84 | IGHV1-18 | 19 | 4.04 |
|  | IGHV3-66 | 1725 | 2.54 | IGHV1-3 | 770 | 4.65 | IGHV3-66 | 18 | 3.82 |
|  | IGHV5-51 | 1590 | 2.34 | IGHV4-59 | 762 | 4.61 | IGHV1-3 | 18 | 3.82 |
|  | IGHV4-59 | 1553 | 2.29 | IGHV3-7 | 696 | 4.21 | IGHV3-7 | 14 | 2.97 |
|  | IGHV3-48 | 1356 | 2 | IGHV3-48 | 679 | 4.1 | IGHV5-51 | 13 | 2.76 |
|  | IGHV4-4 | 1260 | 1.85 | IGHV5-51 | 640 | 3.87 | IGHV3-74 | 13 | 2.76 |
|  | IGHV7-4-1 | 1185 | 1.74 | IGHV3-9 | 548 | 3.31 | IGHV4-59 | 12 | 2.55 |
|  | IGHV3-7 | 1136 | 1.67 | IGHV3-21 | 519 | 3.14 | IGHV7-4-1 | 10 | 2.12 |
|  | IGHV3-21 | 1104 | 1.62 | IGHV4-4 | 499 | 3.01 | IGHV3-9 | 10 | 2.12 |
|  | IGHV3-9 | 1060 | 1.56 | IGHV4-34 | 423 | 2.55 | IGHV4-4 | 9 | 1.91 |
|  | IGHV3-15 | 1011 | 1.49 | IGHV3-74 | 397 | 2.4 | IGHV4-39 | 8 | 1.7 |
|  | IGHV3-11 | 917 | 1.35 | IGHV3-11 | 392 | 2.37 | IGHV4-34 | 8 | 1.7 |
|  | IGHV4-31 | 894 | 1.31 | IGHV7-4-1 | 357 | 2.16 | IGHV2-70 | 8 | 1.7 |
| LIGHT CHAIN | Gene | #Sequences | Percentage | Gene | #Families | Percentage | Gene | #Sequences | Percentage |
|  | IGKV1-39 | 6709 | 14.63 | IGKV1-39 | 2123 | 12.84 | IGKV1-39 | 70 | 18.42 |
|  | IGKV3-20 | 3882 | 8.47 | IGKV3-11 | 1504 | 9.1 | IGKV3-11 | 48 | 12.63 |
|  | IGLV1-51 | 2997 | 6.53 | IGKV3-20 | 1335 | 8.07 | IGKV3-20 | 35 | 9.21 |
|  | IGKV3-11 | 2753 | 6 | IGKV4-1 | 1069 | 6.46 | IGKV4-1 | 23 | 6.05 |
|  | IGKV4-1 | 2484 | 5.42 | IGKV1-33 | 789 | 4.77 | IGKV1-16 | 19 | 5 |
|  | IGKV3-15 | 1811 | 3.95 | IGKV2-28 | 777 | 4.7 | IGKV1-33 | 18 | 4.73 |
|  | IGKV1-5 | 1722 | 3.75 | IGKV1-16 | 690 | 4.17 | IGKV3-15 | 15 | 3.94 |

|  |  |  |  |  |  |  |  |  |
| --- | --- | --- | --- | --- | --- | --- | --- | --- |
| IGKV1-12 | 1627 | 3.55 | IGKV1-5 | 669 | 4.04 | IGKV1-12 | 12 | 3.15 |
| IGKV1-33 | 1532 | 3.34 | IGKV1-12 | 669 | 4.04 | IGKV1-5 | 11 | 2.89 |
| IGLV3-19 | 1479 | 3.22 | IGKV3-15 | 633 | 3.83 | IGLV1-40 | 10 | 2.63 |
| IGKV2-28 | 1427 | 3.11 | IGLV2-14 | 561 | 3.39 | IGKV2-30 | 9 | 2.36 |
| IGLV1-47 | 1377 | 3 | IGKV1-27 | 558 | 3.37 | IGKV2-29 | 9 | 2.36 |
| IGLV1-44 | 1367 | 2.98 | IGLV3-1 | 454 | 2.74 | IGKV1-13 | 9 | 2.36 |
| IGLV2-14 | 1310 | 2.85 | IGLV1-44 | 427 | 2.58 | IGLV3-21 | 8 | 2.1 |
| IGLV3-1 | 1264 | 2.75 | IGKV2-30 | 418 | 2.52 | IGKV2-28 | 8 | 2.1 |
| IGKV1-17 | 1123 | 2.45 | IGLV3-21 | 412 | 2.49 | IGKV1-27 | 7 | 1.84 |
| IGLV1-40 | 1089 | 2.37 | IGLV1-47 | 409 | 2.47 | IGKV1-17 | 7 | 1.84 |
| IGKV1-16 | 1076 | 2.34 | IGLV1-40 | 407 | 2.46 | IGLV1-47 | 6 | 1.57 |
| IGLV3-21 | 1007 | 2.19 | IGLV3-19 | 371 | 2.24 | IGKV1-NL1 | 6 | 1.57 |
| IGKV2-30 | 812 | 1.77 | IGKV1-17 | 370 | 2.23 | IGLV3-19 | 5 | 1.31 |

The top heavy and light V region genes are identical among medicinal patented sequences, medicinal patents and therapeutics. The most used human heavy chain V-gene by sequence, family and therapeutic usage is IGHV3-23, accounting for 25.29% of all patented medicinal sequences, occurs in 15.56% of all medicinal families and accounts for 16.38% of therapeutics. The most frequently observed human light chain germline usage is IGKV1-39, accounting for 14.63% of all patented medicinal sequences, 12.84% of all medicinal patent families and 18.42% of therapeutic antibodies. Some of the most commonly observed genes might be the result of specific platform choices^26^ that might attempt to recapitulate naturally observed frequencies^8^ or focus on a small set of scaffolds^27^. The most frequently used germlines are broadly corresponding between patented sequences, medicinal patents and therapeutics, even though the ordering might not be the same. This indicates that the patent literature well reflects the choices of V-region genes of therapeutic antibodies in clinical use.

### Antibodies from patent documents well reflect therapeutic antibody sequences, with the exception of CDR-H3 lengths

The germline gene distribution of antibody sequences from patents appears to reflect the germline gene distribution of therapeutic sequences, though such comparison is not fit to indicate the actual sequence discrepancies between the two datasets. We checked to what extent patented sequences are a reflection of therapeutics by pairwise sequence comparisons between the two datasets.

For each of the 563 therapeutics we checked if we can find a perfect length-matched hit in PAD. For 546 (96.98%) out of 563 therapeutics we found a perfect length-matched hit in PAD. For the remaining 17 therapeutics without perfect matches, we found that the PAD version used for this study (Jan 2020) was out of date or there existed only high sequence identity matches as compared to Lens.org but not perfect ones (Supplementary Table 4).

For each antibody sequence from a patent, we noted the highest IMGT sequence identity to any therapeutic and present the results stratified by AllPatAb and MedPatAb sequences in Figure 3. Large proportion of PAD sequences align with high sequence identity to one of the 563 therapeutics. Total of 21,772 (16.08%) of heavy chain AllPatAb sequences and 17,378 (18,67%) of heavy chain MedPatAb sequences have matches of 90% sequence identity or better to a therapeutic sequence. Total of 44,919 (40,94%) of light chain AllPatAb sequences and 31,241 (46,16%) of light chain MedPatAb sequences have matches of 90% sequence identity or better to a therapeutic sequence. Altogether this illustrates that many sequences in patent documents well reflect the therapeutic antibody sequences currently in the clinical use. However there is also a large number of sequences with matches below 90% sequence identity to either heavy or light therapeutic heavy chain. This could reflect sequences that are only currently in development or never found their way to the clinic as a result of failure, abandonment or otherwise.

**Figure 3.**
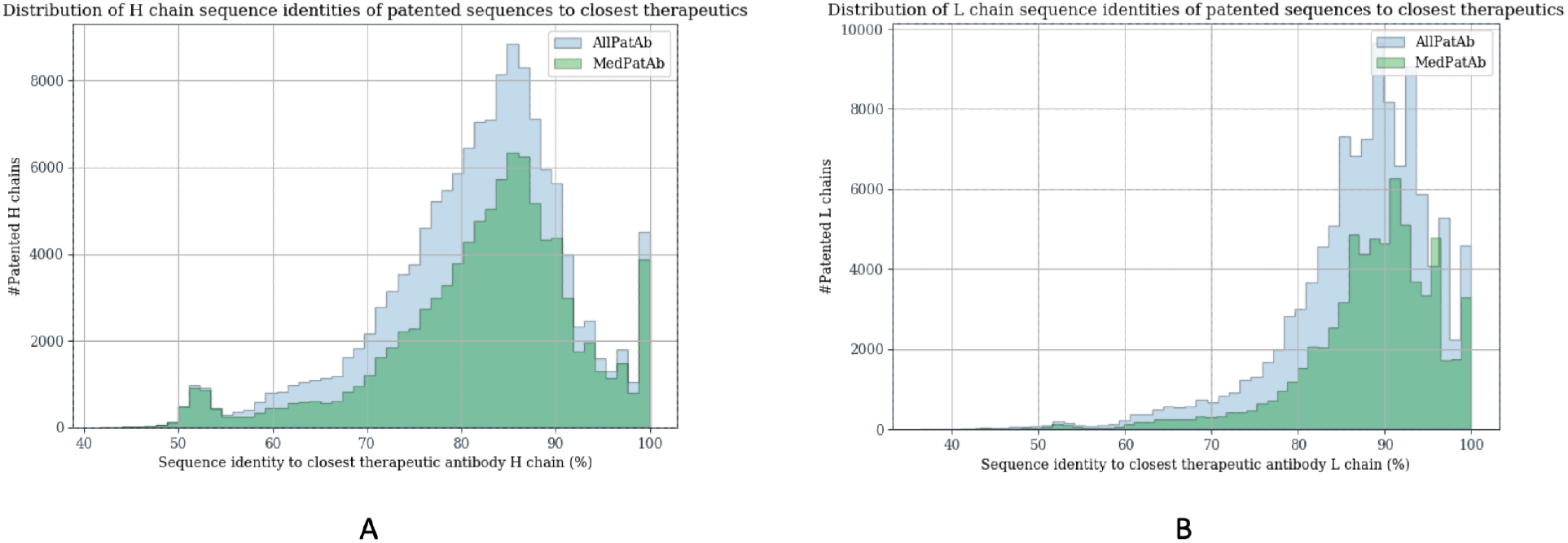
Closest matches of antibody sequences from patents to therapeutic antibodies. For each sequence in AllPatAb and AllPatMed we noted the closest IMGT sequence identity to a therapeutic antibody. A) Distribution of heavy chain sequence identities to closest therapeutic heavy chain. B) Distribution of light chain sequence identities to closest therapeutic light chain.

Perfect matches between full variable region PAD sequences and therapeutics implicitly indicates good correspondence in the CDR region. Arguably, the most diverse and thus the most engineered portion of an antibody is its heavy chain CDR3 region, CDR-H3^29,30^. The length of CDR-H3 has been previously shown to be a good estimator of overall developability of an antibody, with therapeutic antibodies having shorter CDR-H3^6^. We contrasted the CDR-H3 lengths found in PAD to those in therapeutic, structural and natural human antibodies. We extracted CDR-H3s from antibody structures found in the Protein Data Bank^31^ that are regularly collected by the Structural Antibody Database^22^ (SAbDab). The natural human antibodies were sourced from a deep Next Generation Sequencing (NGS) study by Briney et al.^8^ downloaded from the Observed Antibody Space database^17^. We found a total of 58,383 unique CDR-H3s in all PAD sequences (AllPatAb), 37,247 unique CDR-H3s in antibodies from medicinal patents (MedPatAb), 422 unique CDR-H3s in therapeutics, 2021 unique CDR-H3s in structures and 73,217,582 unique CDR-H3s from natural human antibodies. We plotted the distribution of lengths for each of these datasets in Figure 4.

**Figure 4.**
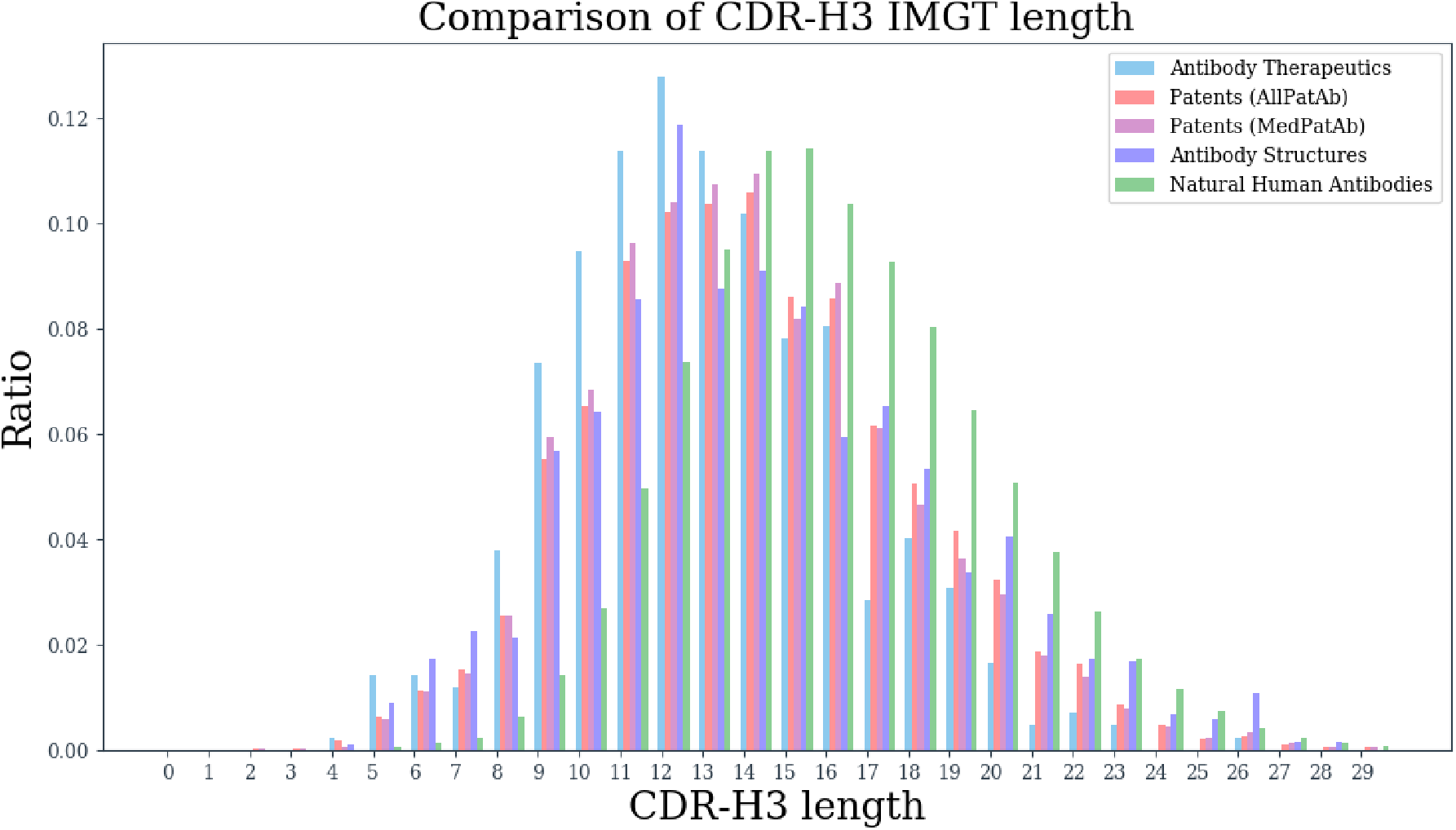
Distribution of CDR-H3 lengths. We plotted the distribution of CDR-H3 lengths from therapeutic antibodies (Antibody Therapeutics), antibodies from patents in PAD (Patents, stratified between AllPatAb and MedPatAb), structures of antibodies from the Protein Data Bank (Antibody Structures) and natural human antibodies from a deep Next Generation Sequencing study (Natural Human Antibodies).

The distribution of CDR-H3 lengths from patent sequences does not appear to be different between AllPatAb and MedPatAb sequences. Therapeutic CDR-H3s have the shortest median lengths, followed by structures, patents and natural human antibodies. The shorter lengths in structures might be reflective of large number of artificial/therapeutic antibodies that can be found in SAbDab^23^. Lengths of CDR-H3s from patent sequences appear to be mid-range between therapeutic and natural antibodies. This suggests that patent antibody sequences might reflect certain amount of engineering of these molecules as they do not follow the natural distribution, normally favoring longer lengths. Nevertheless patent antibody CDR-H3 do not recapitulate the therapeutic CDR-H3 length distribution. Since vast majority of therapeutic CDRs can be found in sequences from patent documents, the discrepancy with the therapeutic length distribution can suggest certain engineering choices faced by those molecules not revealed in this study.

### Patent landscape of single domain antibodies

Our earlier results revealed that majority of antibodies from patents align well to human or mouse germline V region genes, which recapitulates the widespread use of ‘traditional’ antibody format containing both heavy and light chains. The third most commonly identified organism was alpaca (Table 4), which suggests the single domain antibody (sdAb) format. The single domain antibodies are found naturally in camelids (camels, lamas, alpacas) and because of the lack of light chain are believed to have more favorable biophysical properties than antibodies, without detriment to their antigen recognition ability^32,33^. They have been commercialized as therapeutics by Ablynx under the protected name Nanobody^®^ with first single domain antibody drug, Caplacizumab, recently approved^34^. Allowing the first sdAb drug in clinical use holds the promise of more molecules in this format in the near future^35^, which can be reflected by patents.

We identified the total number of patent families in PAD having sdAbs to quantify the possible number of molecules in this format in development, providing an orthogonal view to currently known therapeutic candidates^35^. Patent families were classified as containing sdAbs if they were classified as C07K2317/569 (Single domain, e.g. dAb, sdAb, VHH, VNAR or nanobody^®^) or C07K2317/22 (from camelids, e.g. camel, llama or dromedary) or if they contained sequences aligning to alpaca sdAb germlines. Using the classification method we identified 845 families and using the alpaca germline method we found 867 families. There was an overlap between the two, resulting in total of 1,176 families identified as containing sdAbs or 7.11% of all of our 16,526 families in PAD. Of the 1,176 families 586 (49.82%) were classified as containing antibodies for therapeutic purposes.

The top 30 organizations sorted by the number of families containing sdAb sequences (Supplementary Table 5) well reflect the companies developing biotherapeutics in this format^35^. The list however contains more organizations than those currently reported as developing sdAb therapies, suggesting that the field might be more nuanced, notwithstanding wide use of sdAbs for imaging and diagnostic purposes^36^. From the list of known sdAb therapeutics, our list does not contain AdAlta and Ossianix that report shark single domain antibodies, sequences of which we do not identify. In fact, not all sdAbs that we identified follow the natural camelid format, as there exist sequences of single domain human antibodies (e.g. US2011097339).

We checked the total number of sequences in PAD that could be identified as sdAbs. The 1,176 patent families that we identified as containing sdAbs hold a total of 48,849 unique heavy chain sequences. Not all of such sequences are sdAbs as the patent document might have included traditional antibodies as well. Therefore we calculated the number of sequences that were identified as alpaca sdAb germlines and sequences found in one of the 1,176 families but containing only heavy chains. We found a total of 12,914 unique sequences aligning to sdAb alpaca germlines and 13,368 unique sequences found in 1,176 sdAb families containing heavy chains only. There was an overlap between the two sequence sets and combining them resulted in a total of 15,792 possible sdAb sequences, which makes up 11,66% of all the 135,397 heavy chain sequences in PAD. Of the 15,792 possible sdAb sequences, 8,342 (52.82%) were found in patent documents classified as containing antibodies for medicinal purposes. Therefore, single domain antibody sequences appear to make up a non-trivial proportion of antibody sequences found in patents which could be indicative of upcoming sdAb clinical trials and approvals.

In order to provide an indication of the possible activity to come in the field of single domain antibodies, we plotted the earliest and most recent dates associated with any of the 586 sdAb patent families classified as having antibodies for medicinal applications (Figure 5). There appears to be a steady increase in the number of patent documents including sdAb sequences for medicinal purposes (same holds true for all 1,176 patent families containing sdAb sequences, Supplementary Figure 2). Given the steady rise in the number of patents containing sdAbs and recent approval of Caplacizumab, one might expect more molecules in this format in clinical use in the future.

**Figure 5.**
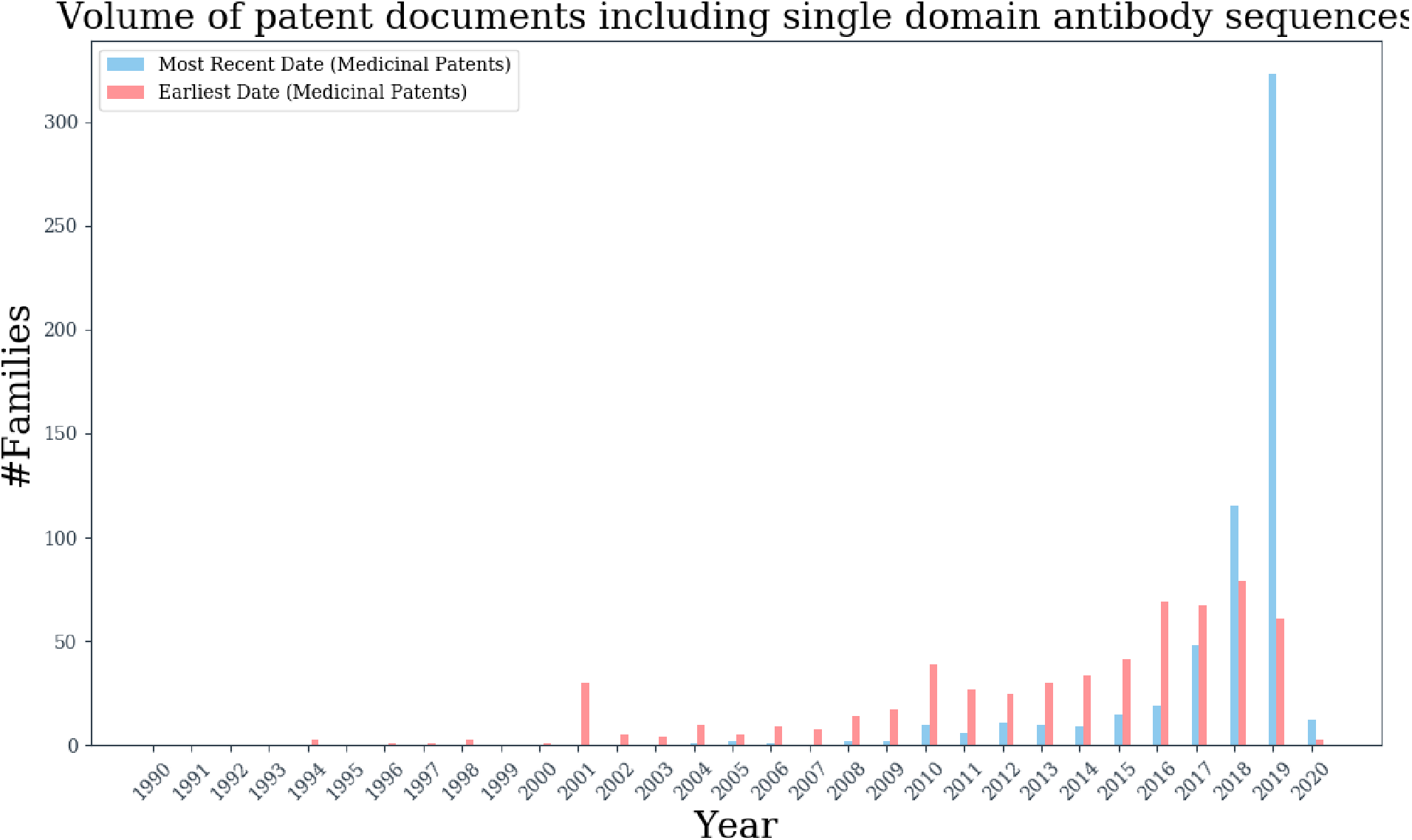
Patents including single domain antibody sequences over time. For each of the 586 patent families in PAD identified as having sdAbs and classified as containing antibodies for medicinal purposes, we noted the earliest and most recent dates, given as red and blue bars respectively.

## Discussion

Successful exploitation of antibodies as therapeutics relies on ever deeper understanding of the biology of these molecules. Many features of therapeutic antibodies can be found in naturally sourced sequences^5^, however effective biotherapeutic requires bespoke engineering for clinical safety and developability^2^. We proposed that such biotherapeutic engineering knowledge could be reflected in patent documents containing antibody sequences.

Our analysis of patents containing antibody sequences revealed that majority of such documents are explicitly developed as containing antibodies for medicinal purposes. Vast majority of therapeutic antibody sequences can be found in patent documents. Further to that, many sequences from patents are within close sequence identity of therapeutic antibodies that are approved or undergoing clinical trials. This suggests that thousands of antibodies from patents could provide a reflection of engineering choices that were made during development therapeutic molecules. Such data could offer an integrated collection of insights into the features that were designed into antibodies to make them successful therapeutics.

This information could be readily exploited by computational methods^4^. It was previously demonstrated that only 137 Clinical Stage Therapeutic (CST) antibodies can provide insights into developability of these molecules^6^. As demonstrated by our analysis, there is an order of magnitude more patented sequences that are close sequence matches to such CSTs. These could indicate different variants and possible features of biotherapeutics, creating a more wholesome picture of what makes a successful biotherapeutic.

Employing antibody sequences from patents however is not without its caveats. Unlike academic literature, patent documents are not designed to convey knowledge but rather offer legal protection. This might result in wide claims on sequence identities to proposed antibody variants that could obfuscate the resulting therapeutic sequence. As we demonstrated, antibodies in patents provide a good reflection of the therapeutics either approved or in clinical use. This would suggest that even though claims could be quite wide on sequence space, many of them appear to fall within the sequence identity orbit of currently available therapeutics. Therefore, certain antibody sequences from patents could broadly reflect the engineering choices in the design of these molecules.

The already large amount of antibodies from patent documents will most likely keep rising, as we demonstrated by the growth in the number of such documents in the recent years. In fact studying such patents could provide an early indication of approvals to come^37^. This might be specifically true in the sphere of single domain antibodies. There is just one such approved therapeutic on the market^34^ and ten in clinical trials^35^ (in 2019). We find a great number of sdAb patents suggesting that the field might further develop in the near future, providing an alternative to traditional monoclonal antibody therapy.

The ongoing increase of patents containing antibodies for medicinal indications will keep contributing to an already ample body of knowledge of antibody engineering. This data could be used to offer insights into the engineering choices in designing these molecules, accelerating delivery of biotherapeutics to the clinic.

## Methods

### Identifying antibody sequences in patent documents

Raw biological sequence data associated with patent documents was downloaded from four freely available accessible services: the United States Patent and Trademark Office (USPTO, https://www.uspto.gov/), the DNA Data Bank of Japan (DDBJ)^12^, European Bioinformatics Institute (EBI)^13^ and World Intellectual Property Organization (WIPO, https://www.wipo.int/). The USPTO data were divided between the full text submissions (https://bulkdata.uspto.gov/) and lengthy sequence listings (http://seqdata.uspto.gov/). Using a custom Python script, the USPTO full text submissions were scanned for nucleotide or amino acid sequences and listings containing these, whereas USPTO PSIPS contained sequence listings only. Using a custom Python script the WIPO FTP documents (ftp://tp.wipo.int/pub/published_pct_sequences) were scanned for nucleotide and amino acid sequences. In both cases of USPTO and WIPO, differences in sequence listing formats from different time periods was accounted for by developing a custom Python parser for each case, transferring all the raw sequences and their associated patent numbers into FASTA format. Data from DDBJ and EBI are available through their ftp services (ftp://ftp.ddbj.nig.ac.jp/ddbj_database and ftp://ftp.ebi.ac.uk/pub/databases respectively) and were readily available in FASTA format.

The nucleotide entries were scanned for antibody sequence by using IGBLAST^16^ as described previously^17^, and their amino acid translations were noted. The raw amino acid sequences were scanned for presence of antibodies using ANARCI^18^. We only kept those amino acid sequences where all three CDR regions and all four framework regions could be identified and that contained only 20 canonical amino acids. This resulted in a dataset of IMGT-numbered amino acid sequences, associated with their patent numbers.

### Patent metadata acquisition and antibody target identification

Different patent numbers can point to the same document, submitted across several jurisdictions, termed ‘patent family’. For each patent number associated with a sequence we identified the patent family by using the Open Patent Services API v. 3.2 (developers.epo.org/ops-v3-2). Using the Open Patent Services API, we downloaded the metadata associated with each family which included: family identifier, title, description, patent numbers with associated dates and applicants.

The patent metadata was used for antibody target identification. Even though there exist certain CPC classifications indicating what the antibody should bind to, we noted that they were not universally present. Therefore we performed manual target annotation, supported by Named Entity Recognition (NER). We applied the GENIA NER^38^ parser to the titles and abstracts of patent families. As with scientific publications titles and abstracts can be expected to reflect the most important content of the document^39^, in particular pertaining to the binding mode of the reported antibody. The resulting annotations accelerated the manual process of annotating each of our patent families with possible targets.

### Web Service

We make the data accessible for academic non-commercial use via web service accessible at http://naturalantibody.com/pad. Users can search for antibody sequences by pasting the amino acids of the variable domains. The input sequence is IMGT-numbered. The sequences in PAD are IMGT-aligned to the input sequence and the top 50 best sequence identity matches are displayed.

