## Supplementary for "Data mining patented antibody sequences"

### Supplementary Information

#### Section 1. Top organizations with patent documents that include antibody sequences

We extracted the institutions that were assigned the patents from our 16,526 patent families to test whether they were organizations chiefly aimed at developing monoclonal antibody therapies. We list the top 100 institutions, sorted by the number of patent families listing antibody sequences with the results given in Supplementary Table 1. We listed the entities as presented in the patent and thus in certain cases organizations that were subsequently acquired can be seen alongside their parent companies at the time of writing. For instance, Genentech appears the be the most prolific of the companies, however it is currently part of Roche. This applies to companies that acquired other antibody format as is the case in Sanofi acquisition of Ablynx. Ablynx specializes in nanobodies and its patent contribution is among the greatest among all the companies according to our data. In total, 9 of the top 10 and 69 of the top 100 institutions are pharmaceutical companies that are involved in development of monoclonal antibodies.

Of the top 100 institutions 26 can be classified as Universities or Research Institutes. Patents by such academic institutions are the effect of their commercialization entities such as YEDA, associated with Weizmann Institute that contributed patents using antibodies for medicinal applications such as JP2006014742. We also find companies such as Canon or Samsung whose core activities are not associated with developing monoclonal antibody therapies. Patents contributed by Canon include documents such as US9675715B2, “Contrast agent for photoacoustic imaging” that fall out of scope of medicinal applications, however they do employ antibodies.

Altogether, strong presence of pharmaceutical companies among the top institutions contributing patents with antibody sequences provide an indication that most documents fall within the scope of their core therapeutic competences.

**Supplementary Table 1**. Organizations ordered by the number of patent families with antibody sequences listed in patent documents. The list contains top 100 organizations sorted by number of patent families where we could identify antibody sequences. The organizations are stratified by the institution type (commercial/academic/government) and by their national headquarters. In certain cases, the company listed on the patent is currently owned by another entity which is listed in the ‘Ownership’ column.

| *Rank* | *#Families* | *Organization* | *Link* | *Ownership* | *Organization Type* | *Country* |
|---|---|---|---|---|---|---|
| *1* | 498 | Genentech | https://www.gene.com | Roche | Commercial | USA |
| *2* | 426 | Roche | https://www.roche.com | | Commercial | Switzerland |
| *3* | 308 | Novartis | https://www.novartis.com | | Commercial | Switzerland |
| *4* | 274 | Amgen | https://www.amgen.com | | Commercial | USA |

| | | | | | | |
|---|---|---|---|---|---|---|
| *5* | 261 | Medimmune | https://medimmune.com | AstraZeneca | Commercial | UK |
| *6* | 241 | Chugai | https://www.chugai-pharm.co.jp | | Commercial | Japan |
| *7* | 226 | Merck USA | https://www.merck.com | | Commercial | USA |
| *8* | 216 | Regeneron | https://www.regeneron.com/ | | Commercial | USA |
| *9* | 215 | Abbvie | https://www.abbvie.com | Abbott spin-off | Commercial | USA |
| *10* | 180 | University of California | https://www.universityofcalifornia.edu/ | | Academic | USA |
| *11* | 168 | Janssen | https://www.janssen.com/ | Jonhnson & Johnson | Commercial | Belgium |
| *12* | 139 | HHS | https://www.hhs.gov/ | | Government | USA |
| *13* | 137 | Pfizer | https://www.pfizer.com/ | | Commercial | USA |
| *14* | 134 | BMS | https://www.bms.com/ | | Commercial | USA |
| *15* | 131 | Samsung | https://www.samsung.com | | Commercial | South Korea |
| *16* | 130 | Kyowa Hakko Kirin | https://www.kyowakirin.com/ | | Commercial | Japan |
| *17* | 129 | University of Pennsylvania | https://home.www.upenn.edu | | Academic | USA |
| *17* | 129 | GSK | https://www.gsk.com/ | | Commercial | UK |
| *18* | 127 | Eli Lily | https://www.lilly.com/ | | Commercial | USA |
| *19* | 126 | Biogen | https://www.biogen.com/ | | Commercial | USA |
| *19* | 126 | Ablynx | https://www.ablynx.com/ | Sanofi | Commercial | Belgium |
| *20* | 120 | University of Texas | https://www.utsystem.edu/ | | Academic | USA |
| *21* | 110 | Sanofi | https://www.sanofi.com/ | | Commercial | France |
| *22* | 106 | UCB | https://www.ucb.com/ | | Commercial | Belgium |
| *23* | 98 | Xencor | https://www.xencor.com/ | | Commercial | USA |
| *24* | 96 | Inserm | https://www.inserm.fr/ | | Government | France |
| *25* | 91 | Boehringer Ingelheim | https://www.boehringer-ingelheim.com/ | | Commercial | Germany |
| *26* | 82 | Memorial Sloan Kettering | https://www.mskcc.org/ | | Academic | USA |
| *26* | 82 | Bayer | https://www.bayer.com/ | | Commercial | Germany |
| *26* | 82 | Abbott Lab | https://www.abbott.com/ | | Commercial | USA |
| *27* | 78 | Scripps | https://www.scripps.edu/about/ | | Academic | USA |
| *28* | 77 | ImmunoGen | https://www.immunogen.com/ | | Commercial | USA |
| *29* | 76 | Novo Nordisk | https://www.novonordisk.com/ | | Commercial | Denmark |
| *30* | 74 | Genmab | http://www.genmab.com/ | | Commercial | Denmark |
| *31* | 73 | Macrogenics | https://www.macrogenics.com/ | | Commercial | USA |
| *32* | 70 | CNRS | http://www.cnrs.fr/ | | Government | France |
| *33* | 66 | Daiichi Sankyo | https://www.daiichisankyo.com/ | | Commercial | Japan |
| *34* | 64 | Innate | https://www.innate-pharma.com/ | | Commercial | France |
| *35* | 60 | Seattle Genetics | https://www.seattlegenetics.com/ | | Commercial | USA |
| *36* | 57 | Wyeth | n/a | Pfizer | Commercial | USA |
| *36* | 57 | Morphosys | https://www.morphosys.com/ | | Commercial | Germany |
| *36* | 57 | Alexion | https://alexion.com/ | | Commercial | USA |
| *37* | 55 | Stanford University | https://www.stanford.edu/ | | Academic | USA |

| | | | | | | |
|---|---|---|---|---|---|---|
| *37* | 55 | Dana-Farber Cancer Institute | https://www.dana-farber.org/ | | Academic | USA |
| *38* | 53 | University of Tokyo | https://www.u-tokyo.ac.jp | | Academic | Japan |
| *38* | 53 | Medarex | n/a | BMS | Commercial | USA |
| *38* | 53 | Dyax | n/a | Shire | Commercial | USA |
| *39* | 52 | Shanghai Cell Therapy Group | http://www.shcell.com/ | | Academic/Commercial | China |
| *40* | 51 | Immunomedics | https://www.immunomedics.com/ | | Commercial | USA |
| *41* | 49 | Jiangsu Hengrui | https://www.hrs.com.cn/hren/about_organize.html | | Commercial | China (PRC) |
| *42* | 48 | Vlaams Instituut voor Biotechnologie | http://www.vib.be/ | | Academic | Belgium |
| *42* | 48 | Merrimack Pharmaceuticals | https://www.merrimack.com/ | | Commercial | USA |
| *42* | 48 | Corixa | defunct | | Commercial | USA |
| *42* | 48 | Alder | https://www.alderbio.com/ | | Commercial | USA |
| *43* | 47 | University of Osaka | https://www.osaka-u.ac.jp | | Academic | Japan |
| *44* | 46 | University of Washington | http://www.washington.edu/ | | Academic | USA |
| *44* | 46 | Oncomed | http://www.oncomed.com/ | | Commercial | USA |
| *44* | 46 | Duke University | https://www.duke.edu/ | | Academic | USA |
| *45* | 45 | Human Genome Sciences | n/a | GSK | Commercial | USA |
| *46* | 44 | Xoma | https://www.xoma.com/ | | Commercial | USA |
| *46* | 44 | Prothena | https://www.prothena.com/ | | Commercial | Ireland |
| *46* | 44 | Pierre Fabre | https://www.pierre-fabre.com/en | | Commercial | France |
| *46* | 44 | National Research Council Canada | https://nrc.canada.ca/en | | Government | Canada |
| *46* | 44 | Kymab | https://www.kymab.com/ | | Commercial | UK |
| *47* | 42 | Philogen | http://www.philogen.com/en/ | | Commercial | Italy |
| *47* | 42 | Juno Therapeutics | https://www.junotherapeutics.com/ | BMS | Commercial | USA |
| *47* | 42 | Baylor University | https://www.baylor.edu/ | | Academic | USA |
| *47* | 42 | A*STAR | https://www.a-star.edu.sg/ | | Academic | Singapore |
| *48* | 40 | Shanghai Hengrui | http://www.shhrp.com/ | | Commercial | China (PRC) |
| *48* | 40 | Schering | n/a | Bayer | Commercial | Germany |
| *48* | 40 | Millenium Pharmaceuticals | http://takedaoncology.com/ | Takeda | Commercial | USA |
| *48* | 40 | Cellectis | http://cellectis.com/ | | Commercial | France |
| *49* | 39 | Sorrento Therapeutics | http://sorrentotherapeutics.com/ | | Commercial | USA |
| *49* | 39 | Canon | https://global.canon/en/ | | Commercial | Japan |
| *50* | 38 | City of Hope National Medical Center | https://www.cityofhope.org/ | | Academic | USA |
| *51* | 36 | Massachussets General Hospital | https://www.massgeneral.org/ | | Academic | USA |
| *51* | 36 | Domantis | n/a | GSK | Commercial | UK |
| *51* | 36 | Academia Sinica | https://www.sinica.edu.tw/en | | Academic | China (Republic of) |

| | | | | | | |
|---|---|---|---|---|---|---|
| *52* | 35 | Medical Research Council UK | https://mrc.ukri.org/ | | Government | UK |
| *53* | 34 | Fred Hutchinson Cancer Research Center | https://fredhutch.org | | Non-Profit | USA |
| *53* | 34 | Crucell | n/a | Jonhnson & Johnson | Commercial | Netherlands |
| *54* | 33 | Imclone | n/a | Eli Lilly | Commercial | USA |
| *54* | 33 | Eureka | https://www.eurekatherapeutics.com/ | | Commercial | USA |
| *55* | 32 | Vanderbilt University | https://www.vanderbilt.edu/ | | Academic | USA |
| *55* | 32 | UCL | https://www.ucl.ac.uk/ | | Academic | UK |
| *55* | 32 | Toray Industries | https://www.toray.com | | Commercial | Japan |
| *55* | 32 | Cytomx | https://cytomx.com/ | | Commercial | USA |
| *56* | 31 | Micromet | n/a | Amgen | Commercial | Germany |
| *56* | 31 | Centocor | n/a | Changed into Janssen | Commercial | USA |
| *57* | 29 | Zymeworks | https://www.zymeworks.com/ | | Commercial | Canada |
| *57* | 29 | Rockefeller University | https://www.rockefeller.edu/ | | Academic | USA |
| *57* | 29 | Novimmune | https://www.novimmune.com/ | Sobi | Commercial | Switzerland |
| *57* | 29 | Astellas | https://www.astellas.com/en/ | | Commercial | Japan |
| *58* | 28 | Rinat Neuroscience | n/a | Pfizer | Commercial | USA |
| *58* | 28 | MIT | http://web.mit.edu/ | | Academic | USA |
| *58* | 28 | LFB | https://www.groupe-lfb.com/en/ | | Commercial | France |
| *59* | 27 | Korea Research Institute of Bioscience | https://www.kribb.re.kr/eng/ | | Academic | South Korea |
| *59* | 27 | Adimab | https://www.adimab.com/ | | Commercial | USA |
| *60* | 26 | Acimune | https://www.acimmune.com/ | | Commercial | Switzerland |
| *61* | 25 | Weizmann Institute | https://www.yedarnd.com/technologies/biotechnology-pharma-and-diagnostics | | Academic | Israel |

#### Section 2.

As an extension of Figure 1 in the main manuscript, we supplemented the data on total patent submissions with only these specifically pertaining to medicinal applications and show these in Supplementary Figure 1.

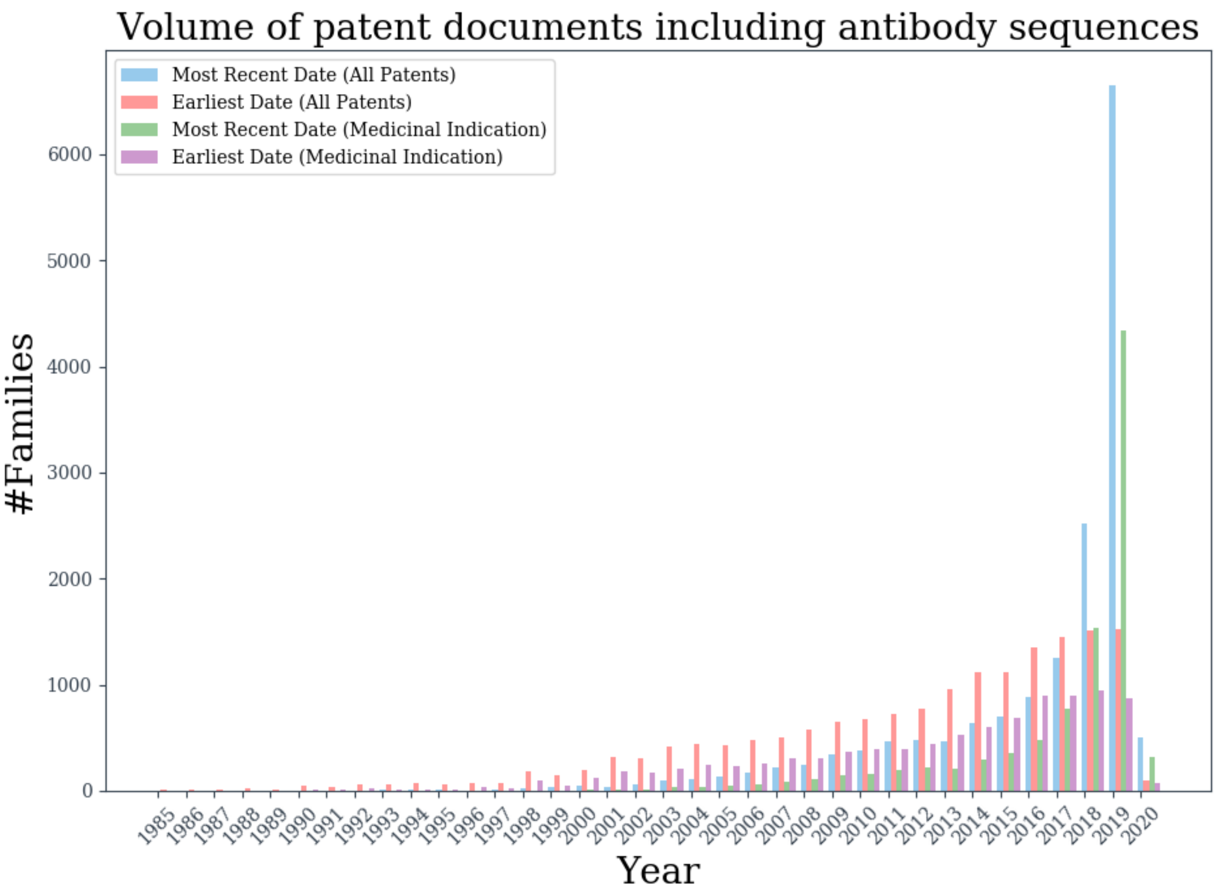


**Supplementary Figure 1.** The volume of patent family documents listing antibody sequences per year. For each patent family we noted the earliest and most recent dates of any documents associated with it and the aggregate numbers of these are given by red and blue bars respectively. We also show the distributions of earliest and most recent dates for the subset of patents classified as including antibodies for medicinal indications, given by purple and green bars respectively.

**Section 3.**

**Supplementary Table 2.** Most common V region gene species antibodies from patents aligned to. Antibodies from all patent documents (AllPatAb) were aligned to fifteen IMGT-derived[1] V region germlines from human, mouse, alpaca, rhesus, rabbit, rat, pig, cow, macaque, zebrafish, trout, salmon, dog, horse and chicken. We noted the number of patent sequences that aligned to the given species germline (#Unique Sequences) and the number of patent families (#Patent Families) these originated from.

| | PER SEQUENCE | | | PER FAMILY | | |
|---|---|---|---|---|---|---|
| **HEAVY CHAIN** | **Organism** | **#Unique Sequences** | **Percentage** | **Organism** | **#Patent Families** | **Percentage** |
| | Human | 94773 | 69.99 | Human | 11287 | 70.29 |
| | Mouse | 21626 | 15.97 | Mouse | 8332 | 51.89 |
| | Alpaca | 12914 | 9.53 | Alpaca | 867 | 5.39 |
| | Rabbit | 2367 | 1.74 | Macaque | 866 | 5.39 |
| | Macaque | 1479 | 1.09 | Horse | 785 | 4.88 |
| | Horse | 1140 | 0.84 | Rabbit | 450 | 2.8 |
| | Chicken | 622 | 0.45 | Chicken | 95 | 0.59 |
| | Dog | 325 | 0.24 | Dog | 43 | 0.26 |
| | Cow | 63 | 0.04 | Rhesus | 32 | 0.19 |
| | Rhesus | 58 | 0.04 | Cow | 27 | 0.16 |
| | Pig | 21 | 0.01 | Pig | 14 | 0.08 |
| | Rat | 3 | 0 | Rat | 5 | 0.03 |
| | Zebrafish | 2 | 0 | Salmon | 3 | 0.01 |
| | Trout | 2 | 0 | Zebrafish | 2 | 0.01 |
| | Salmon | 2 | 0 | Trout | 1 | 0 |
| **LIGHT CHAIN** | **Organism** | **#Unique Sequences** | **Percentage** | **Organism** | **#Families** | **Percentage** |
| | Human | 77965 | 71.06 | Human | 9914 | 64.55 |
| | Mouse | 19258 | 17.55 | Mouse | 8368 | 54.48 |
| | Rhesus | 7806 | 7.11 | Rhesus | 3692 | 24.04 |
| | Rabbit | 2473 | 2.25 | Rat | 592 | 3.85 |
| | Rat | 957 | 0.87 | Rabbit | 442 | 2.87 |
| | Chicken | 718 | 0.65 | Chicken | 99 | 0.64 |
| | Cow | 264 | 0.24 | Cow | 64 | 0.41 |
| | Dog | 228 | 0.2 | Dog | 29 | 0.18 |
| | Pig | 24 | 0.02 | Pig | 14 | 0.09 |
| | Horse | 19 | 0.01 | Horse | 11 | 0.07 |

**Supplementary Table 3**. Top-20 most common human V region genes antibodies from patents aligned to. For each patent antibody sequence in PAD (AllPatAb) that aligned to human germline V regions, we noted the IMGT V region gene. We show the number of unique sequences that aligned to a given human V region gene (#Sequences) and number of patent families these originated from (#Families). We also show the number of therapeutic antibody sequences in clinical use that align to the given V region gene (Per Therapeutic).

| | PER SEQUENCE | | | PER FAMILY | | | PER THERAPEUTIC | | |
|---|---|---|---|---|---|---|---|---|---|
| **HEAVY CHAIN** | **Gene** | **#Sequences** | **Percentage** | **Gene** | **#Families** | **Percentage** | **Gene** | **#Sequences** | **Percentage** |
| | IGHV3-23 | 24181 | 25.51 | IGHV3-23 | 4120 | 24.93 | IGHV3-23 | 77 | 16.38 |
| | IGHV1-2 | 7196 | 7.59 | IGHV1-69 | 2103 | 12.72 | IGHV1-69 | 39 | 8.29 |
| | IGHV1-69 | 7117 | 7.5 | IGHV3-30 | 2058 | 12.45 | IGHV1-46 | 38 | 8.08 |
| | IGHV3-30 | 6542 | 6.9 | IGHV3-66 | 1776 | 10.74 | IGHV3-33 | 26 | 5.53 |
| | IGHV1-46 | 4913 | 5.18 | IGHV1-46 | 1718 | 10.39 | IGHV3-48 | 21 | 4.46 |
| | IGHV1-18 | 3179 | 3.35 | IGHV1-2 | 1634 | 9.88 | IGHV3-30 | 21 | 4.46 |
| | IGHV3-33 | 3041 | 3.2 | IGHV3-33 | 1357 | 8.21 | IGHV1-2 | 21 | 4.46 |
| | IGHV3-66 | 2862 | 3.01 | IGHV1-3 | 1199 | 7.25 | IGHV1-18 | 19 | 4.04 |
| | IGHV1-3 | 2419 | 2.55 | IGHV4-59 | 1197 | 7.24 | IGHV3-66 | 18 | 3.82 |
| | IGHV4-59 | 2227 | 2.34 | IGHV1-18 | 1176 | 7.11 | IGHV1-3 | 18 | 3.82 |
| | IGHV5-51 | 2173 | 2.29 | IGHV3-7 | 1058 | 6.4 | IGHV3-7 | 14 | 2.97 |
| | IGHV3-7 | 2172 | 2.29 | IGHV3-48 | 1053 | 6.37 | IGHV5-51 | 13 | 2.76 |
| | IGHV7-4-1 | 2004 | 2.11 | IGHV5-51 | 996 | 6.02 | IGHV3-74 | 13 | 2.76 |
| | IGHV3-48 | 1934 | 2.04 | IGHV3-9 | 980 | 5.93 | IGHV4-59 | 12 | 2.55 |
| | IGHV3-9 | 1855 | 1.95 | IGHV3-21 | 819 | 4.95 | IGHV7-4-1 | 10 | 2.12 |
| | IGHV4-4 | 1771 | 1.86 | IGHV4-4 | 768 | 4.64 | IGHV3-9 | 10 | 2.12 |
| | IGHV3-21 | 1601 | 1.68 | IGHV4-34 | 687 | 4.15 | IGHV4-4 | 9 | 1.91 |
| | IGHV3-11 | 1404 | 1.48 | IGHV3-11 | 652 | 3.94 | IGHV4-39 | 8 | 1.7 |
| | IGHV3-15 | 1348 | 1.42 | IGHV3-74 | 640 | 3.87 | IGHV4-34 | 8 | 1.7 |
| | IGHV4-34 | 1254 | 1.32 | IGHV7-4-1 | 554 | 3.35 | IGHV2-70 | 8 | 1.7 |
| **LIGHT CHAIN** | **Gene** | **#Sequences** | **Percentage** | **Gene** | **#Families** | **Percentage** | **Gene** | **#Sequences** | **Percentage** |
| | IGKV1-39 | 10544 | 13.52 | IGKV1-39 | 3331 | 20.15 | IGKV1-39 | 70 | 18.42 |
| | IGKV3-20 | 6402 | 8.21 | IGKV3-11 | 2216 | 13.4 | IGKV3-11 | 48 | 12.63 |
| | IGKV3-11 | 5183 | 6.64 | IGKV3-20 | 2139 | 12.94 | IGKV3-20 | 35 | 9.21 |
| | IGKV4-1 | 4625 | 5.93 | IGKV4-1 | 1644 | 9.94 | IGKV4-1 | 23 | 6.05 |
| | IGKV3-15 | 3833 | 4.91 | IGKV1-33 | 1250 | 7.56 | IGKV1-16 | 19 | 5 |
| | IGLV2-14 | 3366 | 4.31 | IGKV2-28 | 1173 | 7.09 | IGKV1-33 | 18 | 4.73 |
| | IGLV1-51 | 3327 | 4.26 | IGKV1-5 | 1091 | 6.6 | IGKV3-15 | 15 | 3.94 |
| | IGKV1-5 | 3074 | 3.94 | IGKV1-16 | 1083 | 6.55 | IGKV1-12 | 12 | 3.15 |
| | IGKV1-33 | 2669 | 3.42 | IGKV1-12 | 995 | 6.02 | IGKV1-5 | 11 | 2.89 |
| | IGKV2-28 | 2493 | 3.19 | IGKV3-15 | 962 | 5.82 | IGLV1-40 | 10 | 2.63 |

| | | | | | | | | |
|---|---|---|---|---|---|---|---|---|
| IGKV1-12 | 2415 | 3.09 | IGKV1-27 | 909 | 5.5 | IGKV2-30 | 9 | 2.36 |
| IGLV1-47 | 2266 | 2.9 | IGLV2-14 | 878 | 5.31 | IGKV2-29 | 9 | 2.36 |
| IGLV3-1 | 2198 | 2.81 | IGLV1-47 | 705 | 4.26 | IGKV1-13 | 9 | 2.36 |
| IGLV1-40 | 2023 | 2.59 | IGLV3-1 | 702 | 4.24 | IGLV3-21 | 8 | 2.1 |
| IGLV1-44 | 2006 | 2.57 | IGLV3-21 | 685 | 4.14 | IGKV2-28 | 8 | 2.1 |
| IGLV3-19 | 1851 | 2.37 | IGLV1-44 | 669 | 4.04 | IGKV1-27 | 7 | 1.84 |
| IGLV3-21 | 1811 | 2.32 | IGLV1-40 | 658 | 3.98 | IGKV1-17 | 7 | 1.84 |
| IGKV2-30 | 1744 | 2.23 | IGLV3-19 | 648 | 3.92 | IGLV1-47 | 6 | 1.57 |
| IGKV1-17 | 1314 | 1.68 | IGKV2-30 | 606 | 3.66 | IGKV1-NL1 | 6 | 1.57 |
| IGKV1-16 | 1286 | 1.64 | IGKV1-13 | 559 | 3.38 | IGLV3-19 | 5 | 1.31 |

**Section 4. Therapeutic sequences without perfect matches to full variable regions in PAD**

For 17 therapeutic antibodies, we did not find a perfect length-matched hit in PAD for either heavy chain, light chain or both. We plotted the best PAD matches to these therapeutics in Supplementary Table 4 and contrasted the results to leading patented sequence search service Lens.org (https://www.lens.org/lens/bio/patseqfinder). The sequence search in Lens.org operates on the basis of blastp[2], searching the Lens repository of sequences mined from patent documents and reports sequence similarity as a result. Therefore wherever there are identical matches, Lens service should be able to identify these if they can be found in their database. In two cases (Icatolimab and Miromavimab) Lens.org identified 100% matches where there were poor matches in PAD. In these two cases, the sequences come from documents that were released after the PAD data for this study was assembled. In all other cases, results from Lens.org are comparable to those in PAD (with the caveat of antibody-specific sequence identity in PAD vs blastp sequence similarity in Lens) and are not 100% matches. This suggests that in certain cases perfect matches cannot be found, but rather very close sequences (~98% sequence identity) that are captured in the patent via stating a certain mutational range from the sequence found in the patent document.

**Supplementary Table 4**. We show the best hits for either heavy or light chain in 17 therapeutics where we could not find a perfect match in PAD (entries for which we could find perfect matches remain empty in the table). For the corresponding light/heavy chains we could not find perfect matches for, we searched Lens.org Patseq Finder to contrast the top results.

| Therapeutic | PAD H best IMGT sequence Identity | PAD L best IMGT sequence Identity | Lens best H sequence blastp similarity (patent number) | Lens best L sequence blastp similarity (patent number) |
|---|---|---|---|---|
| **Abelacimab** | 98.36% | 93.63% | 98.4% (US 2017/0022292 A1) | 98.2% (US 2017/0022292 A1) |
| **Abrezekimab** | 95.83% | 94.39% | 98.3%, (US 2013/0108622 A1) | 98.1% (US 2010/0260773 A1) |
| **Bedinvetmab** | | 99.09% | | 99.1%, (US 2019/0276548 A1) |

| | | | | |
|---|---|---|---|---|
| **Clervonafusp** | | 92.79% | | 94.6% (US 2017/0174790 A1) |
| **Disitamab** | 84.40% | 91.58% | 82.1% (US 2017/0166642 A1) | 92.5% (US 2020/0101142 A1) |
| **Ezabenlimab** | 99.16% | 99.09% | 99.2% (US 2017/0334995 A1) | 99.1% (US 2017/0334995 A1) |
| **Glenzocimab** | | 99.10% | | 99.2%, (US 2018/0236071 A1) |
| **Ianalumab** | | 99.07% | | 99.1%, (US 2010/0021452 A1) |
| **Icatolimab** | 83.20% | 96.42% | 100% (WO 2020/024897 A1) | 100% (WO 2020/024897 A1) |
| **Lecanemab** | | 98.21% | | 98.2% (US 2019/0107537 A1) |
| **Miromavimab** | 90.99% | 98.07 | 100% (WO 2020/089742 A1) | 100% (WO 2020/089742 A1) |
| **Tilavonemab** | | 99.09% | | 99.2% (US 2019/0224339 A1) |
| **Tilogotamab** | | 99.05% | | 99.1% (US 2015/0139997 A1) |
| **Tilvestamab** | 94.91% | | 99.2% (KR 20190021244 A) | |
| **Tomaralimab** | 99.15% | | 99.2% (WO 2019/195409 A1) | |
| **Vilobelimab** | 86.20% | 98.19% | 85.7% (JP 2007202443 A) | 98.2% (US 2013/0243795 A1) |
| **Zuberitamab** | 99.15% | | 99.2% (US 2019/0153471 A1) | |

#### Section 4. Top 30 organizations by patent families containing single domain antibodies.

**Supplementary Table 5.** Top 30 organizations by number of patent families containing single domain antibody sequences.

| #PATENT FAMILIES | COMPANY |
|---|---|
| **128** | Ablynx |
| **45** | Vlaams Instituut voor Biotechnologie |
| **35** | National Research Council Canada |
| **33** | Domantis |
| **31** | GSK |
| **24** | CNRS |
| **21** | Vrije Universiteit Brussel |
| **20** | Ghent University |
| **18** | Boehringer Ingelheim |
| **18** | Amgen |
| **17** | Inserm |
| **16** | University of California |
| **13** | Université Libre de Bruxelles |
| **13** | Roche |
| **13** | Orionis Biosciences |
| **13** | BMS |
| **12** | Novartis |
| **11** | UCB |
| **11** | HHS |

| | |
|---|---|
| **11** | Genentech |
| **10** | voyagertherapeutics |
| **10** | moleculartemplates |
| **10** | Merck USA |
| **10** | Institut Pasteur |
| **10** | Chugai |
| **9** | vhsquared |
| **9** | nanjinglegendbiotech |
| **9** | crescendobiologics |
| **9** | argenx |
| **8** | harpoontherapeutics |

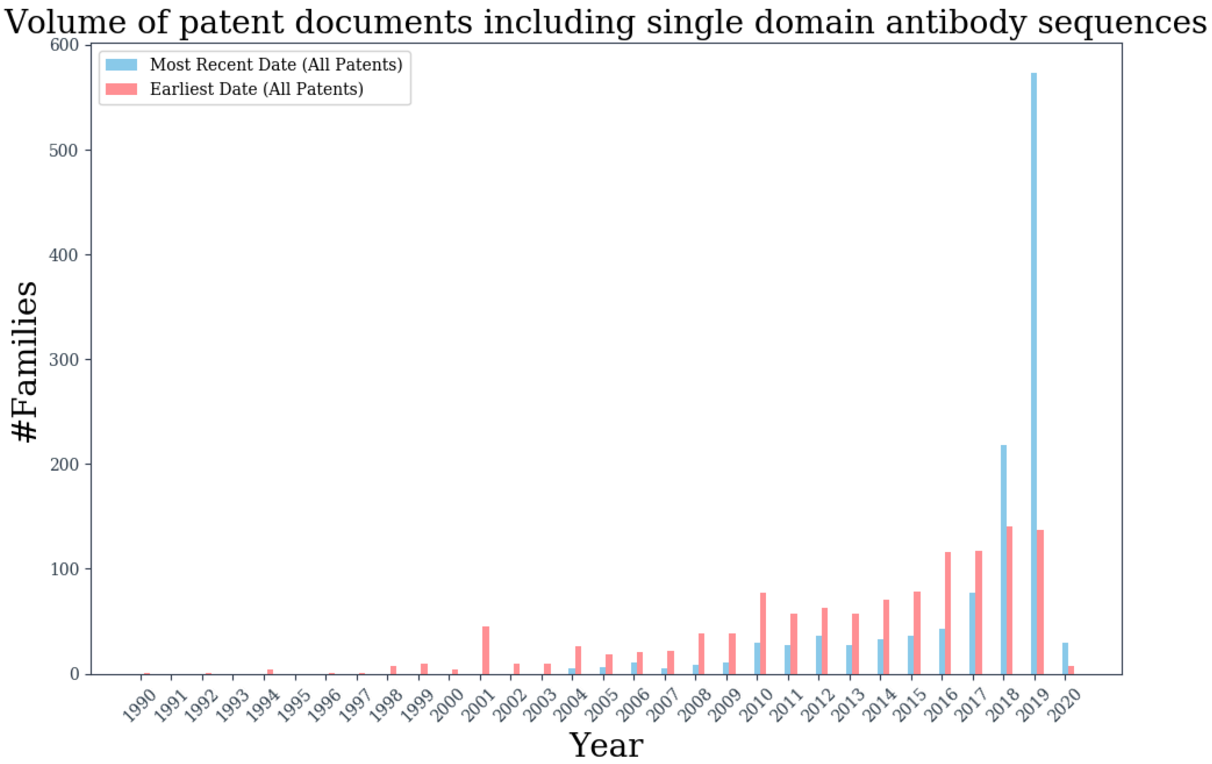


**Supplementary Figure 2.** Patents including single domain antibody sequences over time. For each of the 1,176 patent families in PAD identified as having sdAbs, we noted the earliest and most recent dates, given as red and blue bars respectively.
